# Mapping CD4^+^ T Cell Landscapes in Glioblastoma Reveals Effectors and Bystanders

**DOI:** 10.1101/2025.09.25.678410

**Authors:** Cameron M. Hill, Anthony Z. Wang, Jose Maldonado, Tanner M. Johanns, Allegra A. Petti, Gavin P. Dunn

## Abstract

The tumor immune microenvironment (TME) has been demonstrated to significantly shape glioblastoma (GBM) progression and therapeutic response, yet the role of CD4^+^ T cells remains incompletely defined. Here, by integrating single-cell RNA, surface-protein, and TCR profiling of 39,788 T cells from 16 high-grade gliomas with five matched blood samples, we mapped 23,550 CD4^+^ T cells across 11 states and revealed predominance of CD4^+^ T cells in primary, but not recurrent, tumors, alongside pronounced tumor–blood discordance. Clonal analyses revealed expansion of cytotoxic effector CD4^+^ T cells within tumors and T_EMRA_ CD4^+^ T cells in blood. A small set of dominant clonotypes (0.2% of unique TCRβ sequences) accounted for >6% of all CD4^+^ T cells across both compartments and were shared across different transcriptional subsets, suggesting diverse transcriptional development stemming from a shared progenitor. In contrast, virally annotated clonotypes were broadly dispersed and largely unexpanded, consistent with bystander populations. Collectively, we investigated the cellular and clonal architecture of human CD4^+^ T cells in GBM and highlight the contrast between PBMC and TIL compartments.

## Introduction

Glioblastoma (GBM) is the most common malignant central nervous system (CNS) cancer in adults^1^. With standard therapy of resection, chemotherapy with temozolomide (TMZ), and radiation, recurrence is almost inevitable and median survival is around fifteen to twenty months^2–4^. Thus, new therapies are desperately needed. Immunoediting, which refers to the ways the immune system can shape and ultimately promote cancer growth, likely contributes to the development and progression of GBM^5–11^. The tumor’s subsequent immune escape and low immunogenicity–in part due to its low mutational burden and concomitant low neoantigen burden–in combination with high levels of tumor immunosuppression and heterogeneity, make GBM especially difficult to treat^12^. As such, immunotherapy^13^, which has been successful in other cancers, has shown limited benefit in GBM. A deeper, high-resolution understanding of the cells comprising the immune compartment is therefore needed to better understand the immune response, refine existing therapeutic approaches, and guide new strategies.

While a number of studies have characterized the functions and phenotypes of CD8^+^ T cells in the GBM tumor microenvironment, CD4^+^ T cells in the GBM ecosystem have been comparatively understudied. Importantly, CD4^+^ T cells are emerging as critical determinants of antitumor immunity and immunotherapy responses^14,15^. Effector CD4^+^ T cells harbor key anti-tumor functions either directly via secretion of IFN-γ and TNF-α as well as through a cytotoxic phenotype^16–18^. In addition, CD4^+^ T cells can effect indirect antitumor properties via activating cytotoxic CD8^+^ cells and facilitating B cell mediated humoral responses^16^. Conversely, regulatory CD4^+^ T cells (T_regs_) can promote cancer growth through several mechanisms, such as IL-10 secretion or CTLA-4 expression, thereby contributing to the immunosuppressive environment by restraining the activity of effector immune cells^16^.

In many cancers, MHC class II molecules–which classically present antigens to activate CD4^+^ T cells–have been demonstrated to play pivotal roles in combating cancer ^19,20^. CD4^+^ T cells may tune GBM patients’ response to immunotherapy, as they have been implicated in both resistance mechanisms and therapeutic efficacy^19,21,22^. For example, after H3K27M neoantigen vaccination, a patient with diffuse midline glioma who entered complete remission mounted a dominant CD4^+^, but not CD8^+^, neoantigen-specific response^23^. One multivalent neoantigen vaccine for GBM primarily induced CD4^+^ T cell responses, despite the vaccine being designed to induce CD8^+^ T cell responses^24^. Other neoantigen vaccines for GBM have similarly demonstrated the ability to provoke CD4^+^ T cell responses ^25–27^. Importantly, in a phase I trial for a vaccine that targeted MHCII-restricted IDH1(R132H) neoantigen in grade 3 and 4 astrocytomas, patients with immune responses to the vaccine had a two-year progression free survival rate of 0.82, and CD4^+^ T cells reactive to IDH1(R132H) was observed in the lesion of a patient who underwent re-resection^28^

Beyond their roles in modulating responses to immunotherapy, CD4^+^ T cells themselves have promising therapeutic applications, as adoptive TIL therapy or as CAR-T cells. In preclinical GBM, IL13Rα2-targeted CAR-T cells derived from CD4^+^ T cells mediated superior and durable antitumor activity compared with those derived from CD8^+^ T cells ^29^, and in another solid tumor model, CAR-T efficacy against cancer hinged on early activation of the CD4^+^ CAR-Ts^30^. These data motivate a focused analysis of CD4^+^ T cells in GBM.

Prior single-cell studies that include HGG CD4⁺ T cells have displayed heterogeneous CD4⁺ states and immunoregulatory axes (KLRB1/CD161, S100A4, and IL-10–mediated myeloid crosstalk), which we extend here with larger multi-omic resolution^31–33^. We examined CD4^+^ T cell phenotypic and clonotypic states by integrating scRNA-seq, CITE-seq, and V(D)J sequencing data from patients with GBM (IDH-wildtype) and astrocytoma, IDH-mutant, grade 4 (G4A). Building upon our previously published CD8^+^-focused analysis of T cells from the same HGG patients^34^, we demonstrate that the majority of TIL in primary, but not recurrent, GBM, are CD4^+^, consistent with prior studies^35,36^. Intratumoral CD4⁺ subsets were enriched for Treg, *GZM K*^+^ (C4), cytotoxic, and effector-memory programs, contrasting with naïve and T_EMRA_ bias in blood. Incorporating V(D)J sequencing (14,438 CD4⁺ cells had V(D)J data, which included 11,534 unique TCRβ sequences), we found clonal expansion tightly linked to state: cytotoxic CD4⁺ T cells were preferentially expanded in tumor, whereas T_EMRA_ expansion localized to blood. A small head of dominant clonotypes (TCR1–TCR24; 0.2% of unique TCRβ) accounted for >6% of all CD4⁺ T cells and spanned multiple effector states within clones, consistent with shared progenitors diversifying under common cues. In contrast, CD4⁺ clonotypes with exact viral annotations were broadly dispersed and largely unexpanded, suggesting bystander populations. Collectively, these findings extend our understanding of the glioma immune landscape by revealing distinct CD4^+^ T cell populations, and they indicate that PBMC-only immunomonitoring can intratumoral CD4⁺ dynamics most relevant to therapy.

## Results

### CD4^+^ T cells predominate in the GBM TIL landscape and express unique transcriptional signatures compared to those in blood

This analysis leveraged previously published single cell data of TIL derived from 16 HGG tumor samples, 13 GBM (11 primary, 2 recurrent) and 3 G4A (1 primary, 2 recurrent), and 5 matched PBMC samples to explore the CD4^+^ T cell compartment. As previously described, TIL were isolated by fluorescence-activated cell sorting, then underwent single cell barcoding and processing before being profiled using scRNA-seq, CITE-seq and V(D)J (Methods, Figure 1A). Of the total 39,788 T cells detected, 23,550 were identified as CD4^+^ (Methods). Across individual samples, the distribution of CD4^+^ versus CD8^+^ T cells varied, with primary HGG tumors generally enriched for CD4^+^ cells, while PBMCs showed a variable CD4:CD8 ratio (Figure 1B). When we examined the T cell ratios in newly diagnosed vs recurrent tumors, newly diagnosed tumors contained a higher proportion of CD4^+^than CD8^+^ T cells, whereas recurrent tumors displayed the opposite pattern, with CD8^+^ cells predominating (Figure 1C). Of the 23,550 CD4^+^ T cells with scRNA-seq data, 19,850 included CITE-seq data and 14,438 included V(D)J data (Table S1).

**Figure 1:**
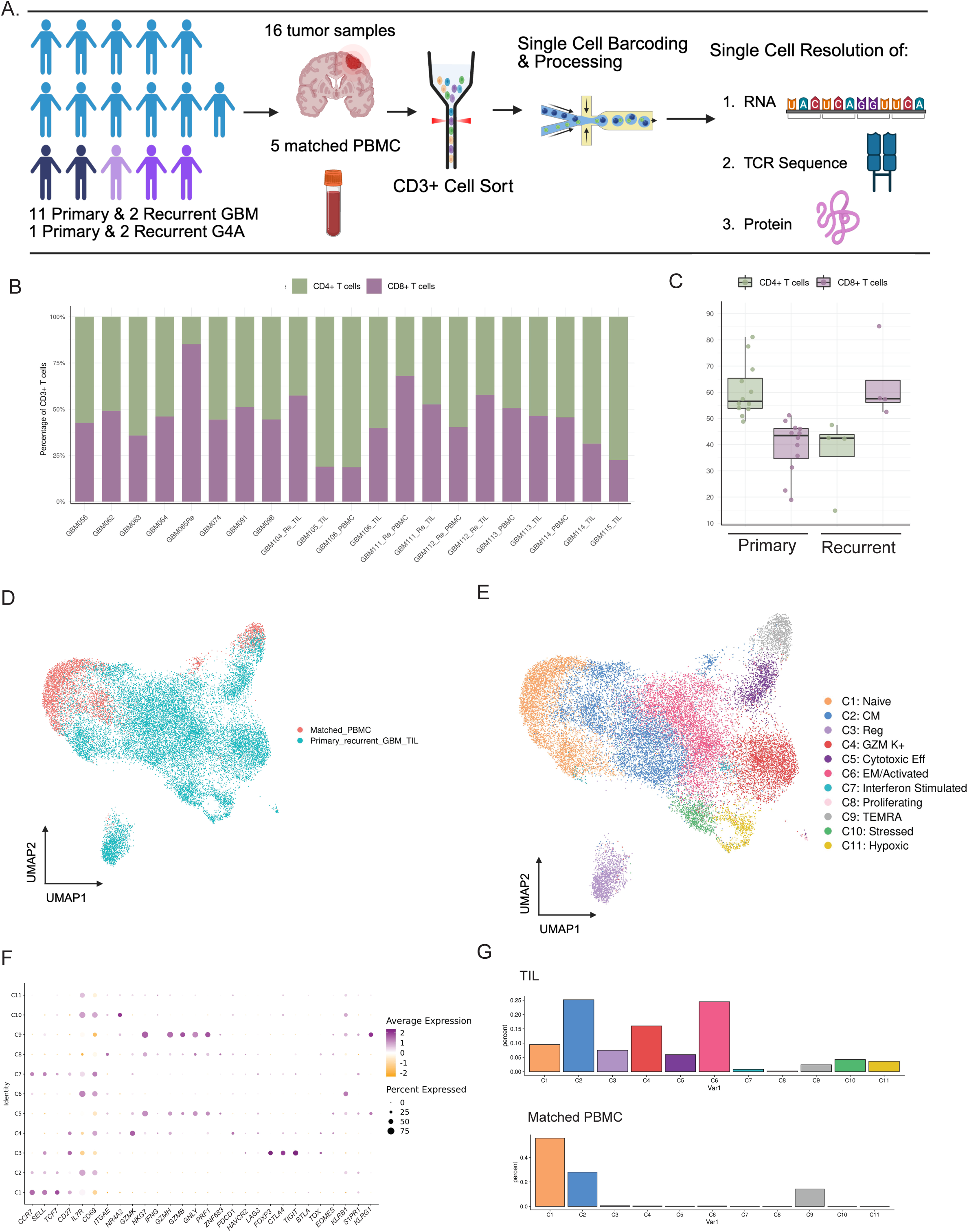
CD4+ T cells predominate in primary glioblastoma tumors and subsets differ between tumor and matched blood. **(A)** Schematic of study design. T cells were isolated from primary and recurrent glioblastoma (GBM) tumors and matched PBMC, followed by scRNA-seq, V(D)J profiling (TCR sequence data) and CITE-seq (protein data). Light blue figures represent patients with primary GBM; dark blue figures represent patients with recurrent GBM; light purple figures represent patients with primary G4A; dark purple figures represent patients with recurrent G4A. **(B)** Distribution of CD4⁺ versus CD8^+^ T cells across individual tumor and PBMC samples. Each bar represents one sample. **(C)** Boxplot showing proportion of CD4⁺ vs CD8⁺ T cells in primary vs recurrent tumors. **(D)** UMAP visualization of T cells colored by sample type (tumor versus PBMC). **(E)** UMAP of all 23,550 CD4⁺ T cells from all HGG and PBMC samples colored by cluster identity (C1–C11; Naive, Central Memory, Regulatory, GZMK⁺, Cytotoxic Effector, Effector Memory/Activated (EM/Activated), Interferon Stimulated, Proliferating, T_EMRA_, Stressed, Hypoxic.**(F)** Dot plot showing RNA expression of select marker genes across CD4⁺ T cell subsets. **(G)** Distribution of CD4⁺ subsets across tumor and blood compartments.

UMAPs colored by compartment showed that TILs clustered apart from PBMC-derived cells, consistent with tumor-specific transcriptional states (Figure 1D). After performing unsupervised clustering and uniform manifold approximation and projection (UMAP) analysis of just the CD4^+^ T cells using joint RNA and protein features, eleven CD4^+^ subsets were identified: Naive, Central Memory, Regulatory, Granzyme K^+^ (*GZM K*^+^), Cytotoxic Effector, Effector Memory, Interferon Stimulated, Proliferating, Terminally Differentiated Effector Memory (T_EMRA_), Stressed, and Hypoxic (Figure 1E, 1F). Subset distribution differed markedly by compartment (Figure 1G) with TIL enriched for T_reg_ (C3), *GZM K*^+^ (C4), Cytotoxic (C5), and Effector Memory (C6) CD4^+^ T cells while PBMC contained higher proportions of naive (C1) and T_EMRA_ (C9) CD4^+^ T cells.

### Clonal expansion is enriched in cytotoxic and T_EMRA_ CD4^+^T cells

Given these differences in subset composition, we were then interested in investigating the clonotypic landscape of CD4+ T cells in GBM. We defined “expanded” as clonotypes detected in ≥3 CD4+ cells. UMAP visualization revealed expanded clonotypes distributed across multiple subsets (Figure 2A). Globally, CD4^+^ TIL contained a higher proportion of expanded clonotypes compared to PBMC (Figure 2B). Repertoire diversity, expressed as normalized TCRβ richness (unique TCRβ per 100 CD4⁺ cells), varied across clusters (Figure 2D). Unsurprisingly, diversity and expansion were inversely related: clusters dominated by expanded clonotypes exhibited low diversity, whereas clusters with high diversity showed little expansion (Figure 2C-D). Expansion concentrated in Cytotoxic (C5) and T_EMRA_ (C9), while as expected, naïve cells were largely unexpanded (Figure 2C). Notably, T_EMRA_ (C9) T cells displayed high levels of expansion in peripheral blood, whereas Cytotoxic (C5) T cells were preferentially expanded in the tumor microenvironment (Figure 2C). Expanded CD4^+^ T cells upregulated effector/cytotoxic genes (e.g., *GZMB*, *PRF1*, *NKG7*) and downregulated naive-associated genes (e.g., *CCR7*, *SELL*) (Figure 2E). Together, these findings demonstrate that CD4^+^ clonal expansion in GBM is tightly linked to functional state.

**Figure 2.**
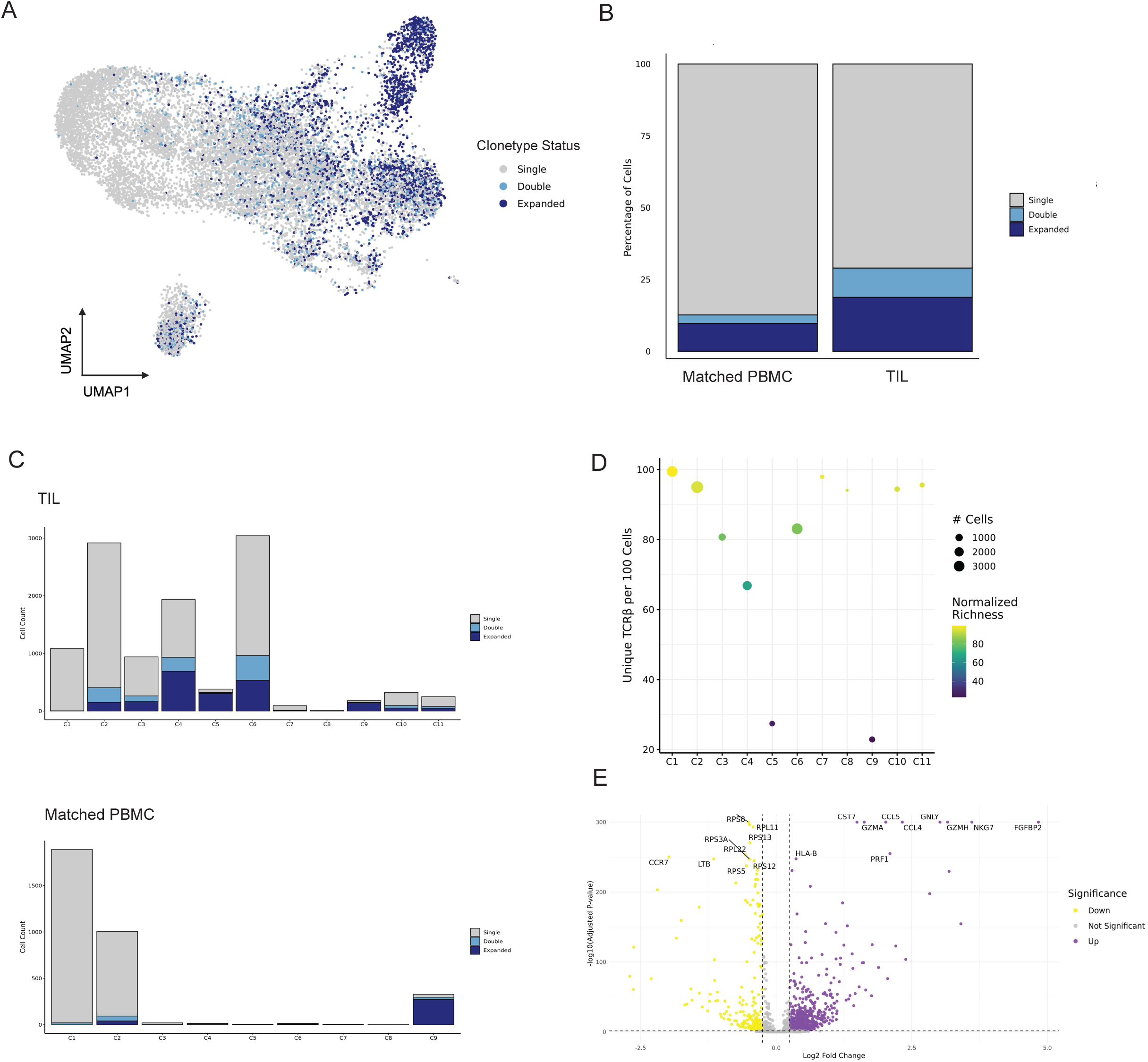
Clonal expansion of CD4+ T cells differs between tumor and blood. **(A)** UMAP of CD4⁺ T cells colored by clonal expansion status (single/double/expanded). **(B)** Proportions of expansion classes (single/double/expanded) by compartment (PBMC vs TIL). **(C)** Distribution of expansion classes within each CD4⁺ cluster, shown separately for TILs and PBMCs **(D)** TCRβ normalized richness (a proxy for diversity) across CD4⁺ T cell clusters **(E)** Differential gene expression in expanded/double CD4⁺ T cells versus singletons (top genes highlighted; thresholds and statistics as specified in Methods).

### A small number of dominant clonotypes account for a disproportionate fraction of the CD4^+^ repertoire

Across 14,438 CD4^+^ T cells with TCR data, there were 11,534 unique TCRβ clonotypes. To discretize the heavy tail-rank frequency distribution while preserving resolution at the head, we ordered clonotypes by abundance and split them into 500 equal size bins (each bin represented ∼0.2% of detected unique clonotypes; 23-24 sequences per group). We chose 500 to yield a manageable size of sequences to visualize individually; coarser groupings (10, 100, 200) produced the same qualitative dominant head (not shown). This analysis highlighted a skewed distribution of clonotypes, with a small subset accounting for a disproportionately large fraction of CD4^+^ T cells, and the steep drop off after the head is consistent with oligoclonal expansion (Fig. 3A). In particular, group one clonotypes (the 24 most frequent; TCR1–TCR24) comprised 0.2% of unique sequences yet represented >6% of all CD4^+^ T cells with TCR data.

**Figure 3.**
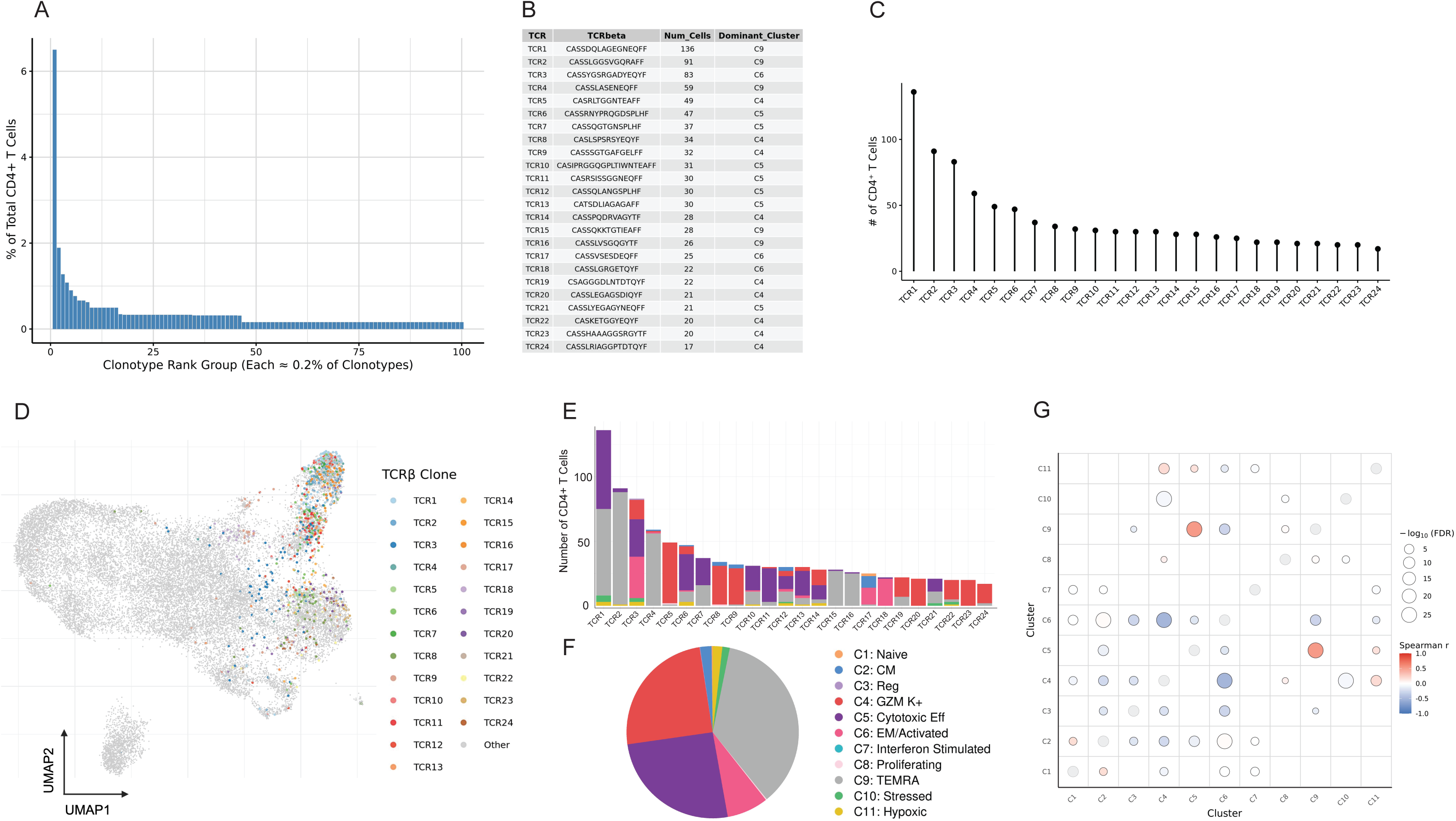
Dominant CD4+ TCRs are distributed across multiple clusters. **(A)** Bar plot of rank bins showing the clonal distribution of TCRβ sequences among CD4⁺ T cells. The x-axis is divided into 500 bins, each representing approximately 0.2% of all unique clonotypes (∼23–24 clonotypes per bin, based on 11,534 unique clonotypes). For clarity, only the first 100 bins are shown. The y-axis indicates the percentage of total CD4⁺ T cells represented in each bin, providing a visual overview of clonal expansion. **(B)** Table summarizing group one clonotypes, the top 24 most expanded TCRβ sequences across the CD4⁺ T cell compartment. For each clonotype, the CDR3β sequence, total number of associated CD4⁺ T cells, and dominant cluster identity are shown. These clonotypes are ranked by frequency and referred to as TCR1 through TCR24 throughout the figure panels. **(C)** Lollipop plot showing the number of CD4⁺ T cells belonging to TCR1–TCR24, arranged in descending order of abundance. **(D)** UMAP highlighting CD4⁺ T cells carrying TCR1–TCR24; each clonotype is color-coded and labeled (all other clonotypes in grey) **(E)** Stacked bar plots showing the cluster (C1–C11) distribution of CD4⁺ T cells expressing each of the top 24 TCRβ clonotypes. **(F)** Pie chart summarizing the overall cluster distribution of all CD4⁺ T cells expressing the top 24 TCRβ clonotypes. **(G)** Spearman correlation matrix of expanded clonotypes (≥3 CD4⁺ cells) co-occurrence across clusters; circle size reflects significance and color encodes Spearman’s ρ; see Methods for details).

TCR1-TCR24 were examined in detail. Their individual clonotypes varied substantially in size, ranging from 17 to 136 and the “dominant cluster” of each clonotype, defined as the cluster with the highest percentage of T cells in the clonotype, belonged to either *GZM K*^+^ (C4), Cytotoxic (C5), EM/Activated (C6), or T_EMRA_ (C9) (Figure 3B-C). TCR1 contained 136 cells, the only clonotype with greater than 100 cells, and had a dominance of T_EMRA_ (C9) T cells (Figure 3B, 3E). UMAP projection highlighted their distribution across phenotypic clusters, revealing that some clonotypes localized tightly to single clusters, like T_EMRA_ (C9) or *GZM K*^+^ (C4), while others were distributed across multiple subsets (Figure 3D). Aggregating across TCR1–TCR24, the majority of cells localized to *GZM K*^+^ (C4), Cytotoxic (C5), and T_EMRA_ (C9), (Figures 3E-F). Finally, correlation analysis of all TCR clonotypes comprised of three T cells or more revealed clusters with highly similar expanded TCR repertoires, particularly those in Cytotoxic (C5) and T_EMRA_ (C9), suggesting expanded T cells derived from shared lineages, likely in response to shared initial antigenic or microenvironmental drivers (Figure 3G). Alternatively, some clusters, like C3 (Tregs) displayed no correlation or anticorrelation from all other clusters, emphasizing their distinct developmental trajectory.

### Viral-specific clonotypes are largely unexpanded and likely represent bystanders

To interrogate the reactivity of the CD4^+^ cells, we next examined CD4^+^ clonotypes with exact CDR3β matches in VDJdb annotated as reactive to common viruses (Influenza A/B, EBV, CMV, SARS-CoV-2/SARS-CoV, HSV-2). Viral TCR+ CD4^+^ T cells were broadly distributed across the UMAP, with representation in multiple clusters rather than confinement to a single state (Figure 4A). Quantification by cluster identity revealed viral-specific clonotypes in nearly all subsets, with relative enrichment in C7 (Interferon Stimulated), and marked under-representation/absence from C8 (Proliferating), T_EMRA_ (C9), and Cytotoxic (C5), which we previously found to be associated with expanded clonotypes (Figure 4B).

**Figure 4.**
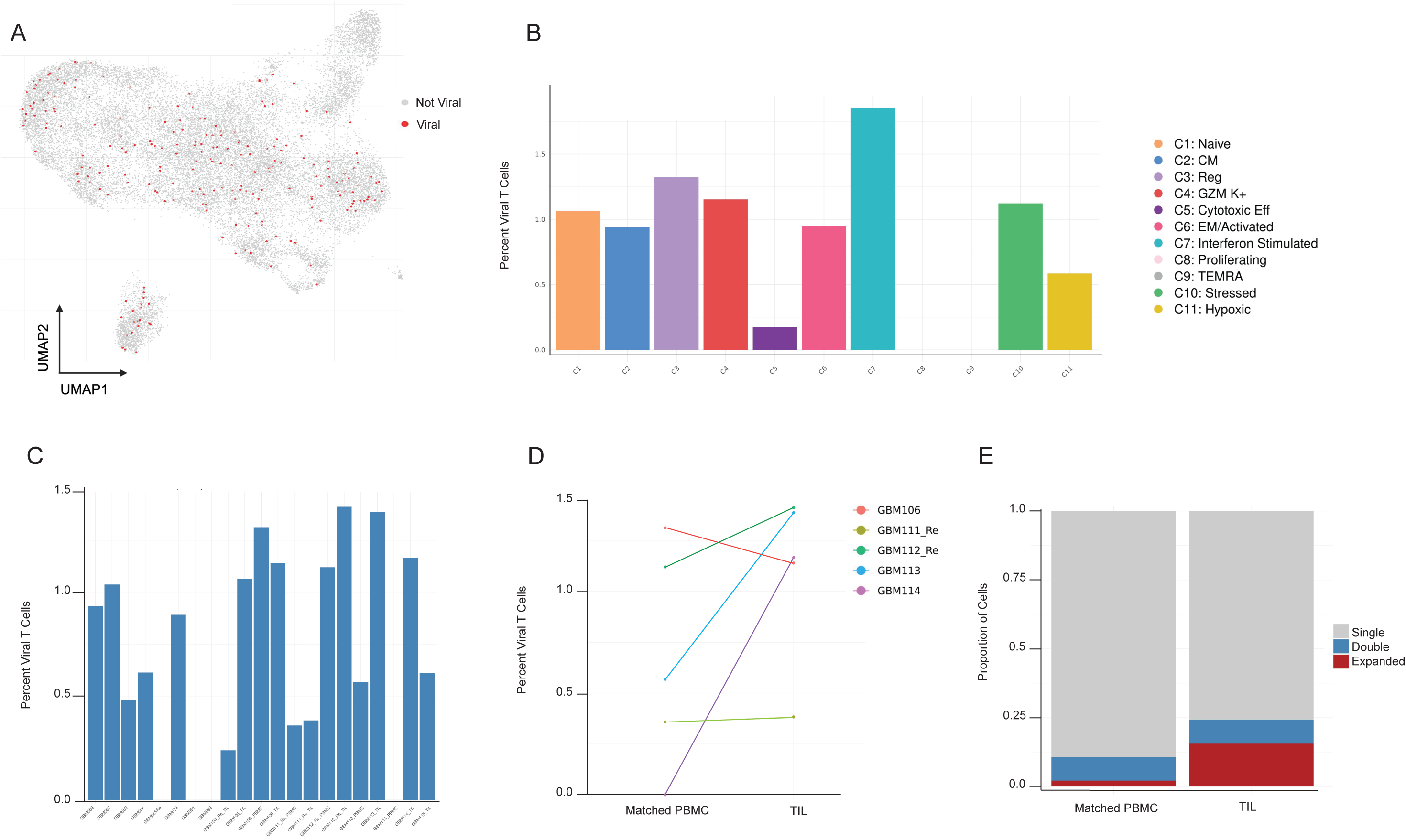
Virus-annotated clonotypes are broadly distributed and minimally expanded in GBM. **(A)** UMAP projection of CD4⁺ T cells highlighting (red) those carrying TCRβ sequences annotated as reactive to common viruses (Influenza A/B, EBV, CMV, SARS-CoV-2/SARS-CoV, HSV-2) based on the VDJdb database. **(B)** Bar plot showing proportion of viral-specific CD4⁺ T cells across clusters (colors = cluster identities) **(C)** Abundance of virus-annotated CD4⁺ T cells across individual patient samples; each bar represents one sample. **(D)** Patient-level comparison of viral-specific TCRs between tumor and peripheral blood compartments. Each line connects matched tumor and blood samples from the same patient. **(E)** Expansion status of viral-specific TCRs in tumor versus blood. Bars indicate the fraction of viral TCRs classified as single, double or expanded within each compartment.

At the sample level, viral-specific TCRs were detected across most patients in both tumor and blood with variable abundance (Figure 4C). In paired tumor–blood analyses, viral clonotypes were usually present in both compartments but at differing frequencies (Figure 4D). Most viral-annotated clonotypes were unexpanded, with only a minority showing expansion (Figure 4E), and the tumor-blood difference was not significant (data not shown). These features are consistent with a bystander phenotype.

## Discussion

In this study, we established what is to our knowledge the largest single-cell, multi-omic atlas of CD4^+^ T cells in HGG. (Prior single-cell GBM studies profiling CD4⁺ T cells, including Mathewson et al., Abdelfattah et al., and Ravi et al., analyzed 8,034, 9,181, and 3,602 CD4⁺ T cells, respectively, all fewer than in our dataset (n = 23,550))^31–33^. Our atlas allowed us to illuminate the landscape of CD4^+^ T cells in HGG based on single cell resolution of RNA, protein, and TCR data and compare their features in tumor vs circulating blood. Our analyses demonstrated that CD4^+^ T cells predominate within the TME of primary HGG and exhibit marked phenotypic divergence compared to circulating CD4^+^ T cells from matched peripheral blood. Previous studies have provided contradictory results on the proportion of T cells that are CD4^+^ versus CD8^+^ in GBM^35–38^. Likely, these contradictions are secondary to the method of T cell quantification, many of which used a limited number of select fields from H&E slides to count T cells and to other confounders (i.e. steroid use, treatment timing, sampling variability). In our study, we demonstrated that the majority of T cells in primary HGG, as assessed by single-cell RNA, protein, or both, are CD4^+^. In other words, we show that the CD8⁺/CD4^+^ ratio is <1, which is consistent with findings reported by Yu et al and Han et al ^35,36^. By contrast, recurrent samples showed CD8^+^/CD4^+^ >1; Sayour et al similarly showed a decrease in the percentage of CD4^+^ T cells in recurrent tumors^39^. Treatment status could underlie this difference, as Ellsworth et al reported that treatment with RT/TMZ results in a decrease in the number of circulating CD4^+^ T cells but not CD8^+^ T cells^40^, and recurrent samples were presumably taken after RT/TMZ treatment. Further work will be necessary to understand the implications of the differing frequencies of CD4⁺ and CD8⁺ T cells in primary vs. recurrent tumors.

Incorporating V(D)J data, we found that clonal expansion was correlated with transcriptional profile. Expanded clonotypes were enriched within Cytotoxic and T_EMRA_ subsets, while naïve-like CD4^+^ T cells remained clonally diverse and unexpanded. This pattern was evident both at the global repertoire level and within dominant individual clonotypes. This supports a model in which antigen exposure within the GBM microenvironment promotes selective CD4^+^ effector differentiation and aligns with reports that human CD4^+^ T cells can acquire direct cytotoxic programs^17,18^.

Presumably such CD4⁺ T cell mediated antitumor function in GBM requires MHC II antigen presentation, provided either by tumor cells or APCs, to prime or enable direct recognition. Malignant glioma tumors have been observed to express MHCII by immunohistochemistry, and malignant glioma cell lines stimulated with IFN-γ have been reported to present nonmutant native peptide to CD4⁺ T cells^41^, establishing that in principle glioma cells can be CD4⁺ T cell targets. In oncolytic HSV, a recent study in murine GBM showed oHSV upregulates MHCII on residual tumor and expands tumor-reactive CD4^+^ T cells, and that CD4⁺ mediated killing of tumor cells is MHCII dependent; in parallel, in an oHSV clinical trial in humans, upregulation of MHCII genes correlated with improved survival^42^. They also found that patients in the Cancer Genome Atlas program with high levels of CD4^+^ T cells and high MHCII expression were reported to have significantly longer median survival than patients with high levels of CD4⁺ T cells and low MHCII expression^42^. Complementary data from another oHSV trial for GBM linked increased clonal diversity of T cells with increased survival, consistent with a polyclonal T cell response in GBM. Finally, evidence that peripheral CD4^+^ memory T cells can recognize patient-specific neoantigens^43^ underscores that CD4^+^ specificity against tumor exists. Together with our observation of expanded, cytotoxic effector CD4⁺ states in human HGG, these data support a model where MHCII in tumors can provide local signal to CD4⁺ T cells that permit direct MHCII dependent tumor recognition and killing. Avenues to amplify this mechanism will have broad therapeutic application.

Notably, a small set of dominant clonotypes (TCR1–TCR24) accounted for a disproportionate share of the CD4^+^ repertoire (representing over 6% of all CD4^+^ T cells despite comprising only 0.2% of unique TCRβ sequences). While some dominant clonotypes localized to one state primarily, like T_EMRA_ or *GZM K*^+^ (C4), the dominant clonotypes that contained cytotoxic subsets were distributed across multiple effector clusters. This “within-clone” heterogeneity is consistent with the idea that individual naïve CD4^+^ T cells can generate heterogeneous effector progeny following antigen encounter, with fate influenced by intrinsic TCR properties, signal strength, and contextual cues^44^. These data favor a framework of shared clonotypes across different subsets in the setting of common upstream cues (antigenic context and/or microenvironmental signals). This contrasts with our previously published findings in CD8^+^ T cells in the same cohort that primarily showed expansion of a *GZM K*^+^ subset, highlighting the different expansion signatures between CD4^+^ and CD8^+^ T cells. Because our data are cross-sectional, we do not infer directionality between the different effector states within a single clone; clone-aware trajectory analyses are necessary to confirm whether one cluster precedes another or represent parallel endpoints.

Together, these data point to selective pressures shaping the CD4^+^ repertoire in GBM and demonstrate clonal focusing within cytotoxic and effector subsets. This has therapeutic implications: in the previously mentioned oHSV trial, survival after treatment was associated with increased clonal diversity^45^, suggesting that broadening repertoire breadth, not only amplifying a few dominant clones, may be advantageous. Additionally, GBM vaccine trials designed to leverage T cell activation routinely test PBMC for peptide reactivity to monitor response to therapy ^24,46–53^. The goal of these therapies is to induce TIL reactive to targeted peptides. Our observation of strong tumor–blood discordance in CD4^+^ clonotype, expansion, and cluster distribution indicates that PBMC-only assays may miss intratumoral dynamics most relevant to efficacy. To our knowledge, we are the first to display single-clonotype-level differences in CD4^+^ TCR expansion between TIL and PBMC in HGG.

By contrast, CD4^+^ clonotypes annotated as viral-specific were broadly distributed across phenotypic states but were overwhelmingly unexpanded. Viral-specific clonotypes were minimal or absent from clusters with the highest degree of expansion, such as cytotoxic and T_EMRA_ subsets. These features suggest that viral-specific clonotypes largely represent bystander populations within the tumor milieu. This mirrors observations in melanoma that non-tumor reactive T cells were enriched for viral reactive TCRs ^54^. Recognizing the presence of these bystanders is important, as these too have the potential to be harnessed as therapy^55^. Distinguishing bystander from tumor-reactive CD4^+^ populations will be essential for designing therapies that broaden productive antitumor breadth.

Additional work is needed to understand the clinical-translational relevance of our findings to date. Specifically, detailed treatment histories were not available for all participants. Therapy such as RT/TMZ and corticosteroids can impact T-cell compartments and could contribute to the primary–recurrent differences we observe. Also, intratumoral sampling sites were not annotated (core versus margin or peritumoral regions). Spatial heterogeneity is a defining feature of HGGs, and regional sampling bias could influence apparent subset composition and expansion enrichment^56^.

Moreover, tumor samples inevitably include intravascular blood, and we cannot perfectly segregate parenchymal tumor-infiltrating lymphocytes from T cells transiting within tumor vessels at the time of biopsy. To partially mitigate this, we analyzed matched PBMCs when available and TIL features should be interpreted in the context of tumor–blood contrasts.

Although the dataset is comparatively large for GBM T cells, matched blood was available for only five cases and recurrent tumors were few, limiting power for stratified analyses. Pooling GBM (IDH-wildtype) and G4A (IDH-mutant, grade 4) increases generalizability across HGG but may blur subtype-specific biology; our study was not powered to dissect subtype-specific CD4⁺ programs. Finally, transcriptional and surface-protein signatures are only surrogates for function: although lack of clonal expansion enrichment suggest a bystander role, functional validation would be required to definitively exclude tumor reactivity. Similarly, a cytotoxic phenotype enriched in the tumor combined with expansion suggests tumor reactivity but does not confirm it. Prospective studies incorporating HLA typing and tumor immunopeptidomics will be required to validate tumor reactivity.

In sum, by assembling the largest single-cell, multi-omic atlas of human CD4⁺ T cells in high-grade glioma, we delineate the cellular and clonal architecture of intratumoral CD4⁺ immunity. We show that CD4⁺ T cells predominate in primary GBM, that tumor and blood contain different phenotypic and clonal landscapes, and that cytotoxic and T_EMRA_ TILs are highly enriched for expansion. A small group of dominant clonotypes (0.2% of unique TCRβ sequences) accounts for >6% of all CD4⁺ T cells with TCR data, and spans multiple effector states, whereas virally annotated clonotypes are broadly dispersed and largely unexpanded, consistent with bystanders. These insights caution against PBMC-only immunomonitoring and motivate strategies to broaden tumor-reactive CD4⁺ breadth.

## Methods

### Study cohort, Sample Collection, and Ethics

All tissue and blood specimens were obtained with written informed consent under IRB-approved protocols as previously described^34^. In brief, adults undergoing resection for high-grade glioma were enrolled; tumors were classified per WHO 2021 (GBM, IDH-wildtype; astrocytoma, IDH-mutant, grade 4 [G4A]).

This study reanalyzed the CD4^+^ T-cell compartment from the same single-cell datasets of cohorts 1 and 2 reported in Wang et al^34^; brain metastasis samples used in that study were excluded here. Sample counts in the present analysis are shown in Fig. 1A (13 GBM, 3 G4A; 12 primary, 4 recurrent; 5 matched PBMCs).

### Single Cell TCR Analysis

Seurat, scRepertoire, and Immunarch were used for clonotype analysis. Clonotype states were defined as the following: single (X = 1), double (X = 2), expanded (X ≥ 3), where X is the number of cells in which the clonotype is detected. We use the terms clusters, subsets, and states interchangeably to describe transcriptionally defined CD4⁺ populations.

### Dominant-clonotype ranking

Across 14,438 CD4^+^ T cells with TCR data, 11,534 unique TCRβ clonotypes were observed. Clonotypes were ordered by frequency and partitioned into 500 rank groups (∼0.2% each). The top rank group (Group 1) comprises the 24 most frequent clonotypes, denoted TCR1–TCR24.

### Correlation of clonotype distributions across clusters

For clonotypes with ≥3 CD4^+^ cells (pooled per patient across tumor ± PBMC), we built a clonotype (rows) × cluster (columns) count matrix; each row was normalized to fractions (sum = 1). We then computed Spearman correlations between cluster-specific vectors of clone fractions and assessed significance with two-sided Spearman tests for each unique cluster pair. P-values were adjusted across all non-redundant pairs using Benjamini–Hochberg FDR, and significant associations were visualized as bubbles with radius proportional to −log₁₀(FDR) and color encoding Spearman r.

### Viral-specific TCR annotation

Viral annotation of TCRs was performed by exact CDR3β amino-acid matching to entries in VDJdb^57^. CDR3β data from TCRs reactive to common viruses (Influenza A/B, EBV, CMV, SARS-CoV-2/SARS-CoV, HSV-2) were downloaded from VDJdb.

### Other Statistical Analyses

The Wilcoxon rank-sum test (two-sided) with Bonferroni correction was used for differential gene analysis. P ≤ 0.05 was considered statistically significant. Normalized richness was computed as unique TCRβ clonotypes per 100 CD4⁺ cells per cluster. Statistical analyses were performed with R or Seurat.

